# In-silico Discovery of Genetic Diversity in *Cucumis sativus* var. hardwickii: A Wild Relative of Cultivated Cucumber

**DOI:** 10.1101/2023.03.11.532174

**Authors:** Suniti Rawat, Prashant Kaushik

**Affiliations:** Department of Biotechnology, Sardar Vallabh Bhai Patel University of Agriculture and Technology, Meerut 250110, Uttar Pradesh; Instituto de Conservación y Mejora de la Agrodiversidad Valenciana, Universitat Politècnica de València, 46022 Valencia, Spain

**Keywords:** *Cucumis sativus* var hardwickii, VEP, Genomic variation, SNP, SNV

## Abstract

Genome-wide sequencing data play an important role in evaluating the genomic level differences between superior and poor-quality crop plants and improving our understanding of molecular association with desired traits. We analyzed the obtained 92,921,066 raw reads from genome-wide resequencing of *Cucumis sativus* var. hardwickii through in-silico approaches and mapped to the reference genome of Cucumis sativus to identify the genome-wide single nucleotide polymorphisms (SNPs) and Single nucleotide variations (SNV). Here, we report 19, 74,213 candidate SNPs including 1,33,468 insertions and 1,43,237 deletions and 75 Indels genome-wide. A total of 2228224 identified variants were classified into four classes including 0.01% sequence alteration, 5.94% insertion, 6.37% deletion and 87.66% SNV respectively. These variations can be a major source of phenotypic diversity and sequence variation within the species. Overall, the discovery of SNPs and genomic variants may help predict the plant response to certain environmental factors and can be utilized to improve crop plants’ economically important traits.

## 1. Introduction

*Cucumis sativus* var. hardwickii is one of the wild relatives of cucumbers. It belongs to the Cucurbitaceae family, primarily found in forests and mountain slopes, at altitudes of 700-2000 m, in China, Myanmar, NE India, Nepal and Thailand. The origin of the cucumber is known to be India, where the wild ancestor *Cucumis sativus* var. hardwickii round, and bitter fruit grows (Lee et al., 2020). It is also found in South India, particularly in the state of Kerala. *Cucumis sativus* var. hardwickii (wild cucumber) is a self-compatible, insect-pollinated taxon with a stable genome (2n = 14) and high reproductive output (85%) as compared to cultivated cucumber (73.3%). It is primarily composed of two commercially available vegetables: cucumber *(Cucumis sativus* L. var. sativus; 2n=2x=14) and melon *(Cucumis melo* L; 2n=2x=24) ( Datar et al., 2013; Golabadi et al., 2012). From India to China, the plant seems to have spread eastward, and from China westward, to Asia Minor, North Africa, and Southern Europe. As such, it represents an extreme of *C. sativus* variation and has the potential to expand the genetic diversity available for commercial cucumber breeding (Naegele et al., 2016).

It possesses a number of advantageous characteristics, including a sequential multiple fruiting habit, multiple lateral branching, and a high overall fruit weight per plant (Staub et al. 1993), dark green-coloured fruits, early maturation, and fruit production during the rainy season. It includes several critical genes for the evolution of cultivated cucumber but is underrepresented in the world’s central gene banks (Behera et al., 2010; Mariod et al., 2017). Genetic markers (morphological, biochemical, and molecular) have been used to characterize the genetic variation found in cucumber (Pandey et al., 2013; Singh et al., 2016). Thus, by analyzing a more robust selection of this species, we can better understand the variation present in this closely related population used in plant improvement programs (Kesh and Kaushik, 2021).

Agricultural lands have been compromised by salination, air pollution, and global warming (Arzani and Ashraf 2016). Studies of single nucleotide polymorphisms (SNPs) and significant advances in the use of genomic methods have created new, quick mapping procedures for identifying features and determining their causes. To expand multidisciplinary science and it is necessary to maintain traditional practices in the field of molecular genetics. Because of their low costs and high performance, microarrays and next-generation sequencing are commonly used (Nadeem et al., 2018). Recent microarrays have been applied to biological samples. Although Northern or quantitative PCR has previously been used in gene studies and the ability to assess a more significant number of genes in a single run improved sensitivity and resolution. The primary goal of next-generation sequencing is to read hundreds of thousands or millions of DNA molecules.

The most common type of DNA sequence variation between alleles is single nucleotide polymorphisms (SNPs), which can be revealed by using advanced technologies for genome analysis. SNPs are single base-pair differences (substitutions or deletions) among chromosome sequences that result from point mutations and account for much of the genetic variation in the genome. The binding affinity between a regulatory protein and a regulatory DNA binding site may change if there is a SNP in that site (Fareed et al., 2013; Zaid et al., 2017). SNP markers have become increasingly popular in plant molecular genetics due to their abundance in the genome and their ease of detection at ultra-high-throughput. Despite these benefits, GBS continues to face challenges due to a lack of quick, reliable data imputation algorithms and powerful computers capable of processing and storing large amounts of sequencing data (Patel et al., 2015). Even in the face of such disadvantages or threats, genomic and other next-generation advances increase the pace and benefits of plant breeding. SNPs are the first step toward unravelling evolutionary and genetic relationships and elucidating agronomic characteristics and plant diseases (Rasheed et al., 2017). Therefore, we performed the genome-wide In-silico mining of SNPs and genomic variants from *Cucumis sativus* var. hardwickii and predicted their effects in detail.

## 2. Material and Methods

### 2.1 Raw reads and Mapping

In-Silico investigation has been done with the raw SRA data available in the public domain at NCBI (National Center for Biotechnology Information). The whole genomic data was used for the SNPs and variants calling using the whole genome sequencing data. Raw reads from the whole genome sequence of *Cucumis sativus* var hardwickii were downloaded from the NCBI, SRA with an accession number SRR2096458usingthe Illumina platform 92,921,066 raw reads were generated. These reads were mapped to the *Cucumis sativus* genome using BWA, and the version of the Cucumber genome was ASM407v2 (var. sativus, cv. Chinese Long, inbred line 9930) (Li and Durbin, 2009).

### 2.2 Genome-wide SNPs calling

BWA-mem (v0.7.17) was used with its default parameters. The vcfstats feature of RTG-tools version 3.12 was used to calculate mapping statistics such as total mapped, unmapped, and paired mapped reads and perform additional analysis on the alignment files. We used UnifiedGenotyper from GATK (v.3.8-0) to call SNPs and indels for variant analysis. We separated the raw variant calling data into SNPs and indels using SelectVariants from GATK (v. 4.0.1.2). Then we applied hard-filtering to these raw SNP calls using VariantFiltration from GATK (v. 4.0.1.2) based on these threshold criteria: MappingQualityRankSum of < −12.5, QUAL < 30, QD < 3.0, FS > 30.0, MQ < 30.0, DP < 200, and genotype-filter-expression DP < 15. We then chose bi-allelic variants using VCFtools (version 0.1.15) ( (DePristo et al., 2011; Danecek et al., 2011).

### 2.3 Variant effect analysis

The mapped data was used for Variant effect prediction using Variant Effect Predictor (VEP) software (version (API), 102) to verify the effect of variants (McLaren et al., 2016). The script for Ensembl Variant Effect Predictor is available at (http://www.ensembl.org/info/docs/tools/vep/script/index.html).

## 3. Results

### 3.1 Genome-wide identification of SNPs

We identified a total 19, 74,213 polymorphic SNPs in *Cucumis sativus* var hardwickii in relation to the reference genome of cucumber with MNPs (0.00), Insertions (1, 33,468), deletions (1,43,237), Indels (75) from all the processed reads and 1,62,673 genotypes were missing (Table 1). Furthermore, the heterozygous/homozygous ratio was also predicted for SNPs (0.20%), insertion (0.99%), deletion (0.85%) with total Het/Hom ratio (0.26%) and not detectable for MNPs and indels (Table 1).

**Table 1.** Summary of SNPs identified in *Cucumis sativus* var. hardwickii.

| Analyzed Factors | Values |
| --- | --- |
| Failed Filters | 0 |
| Passed Filters | 6404727 |
| SNPs | 1974213 |
| MNPs | 0 |
| Insertions | 133468 |
| Deletions | 143237 |
| Indels | 75 |
| Same as reference | 3991061 |
| Missing Genotype | 162673 |
| SNP Transitions/Transversions | 1.53 (2189498/1432901) |
| Total Het/Hom ratio | 0.26 (459863/1791130) |
| SNP Het/Hom ratio | 0.20 (327665/1646548) |
| MNP Het/Hom ratio | - (0/0) |
| Insertion Het/Hom ratio | 0.99 (66254/67214) |
| Deletion Het/Hom ratio | 0.85 (65869/77368) |
| Indel Het/Hom ratio | - (75/0) |
| Insertion/Deletion ratio | 0.93 (133468/143237) |
| Indel/SNP+MNP ratio | 0.14 (276780/1974213) |

### 3.2. Genome-wide Variant analysis using Ensembl Variant Effect Predictor

VEP analysis was performed to determine the effect of variants on genes, mRNA transcripts, protein-coding, and regulatory sequences that may help evaluate the genetic diversity within the species.

#### 3.2.1 General statistics data description of analyzed variants

A total of 22,28,224 variants were processed out of 6404727 input reads along with 23,777 number of overlapped genes and similar for overlapped transcripts with zero novel variants and overlapped regulatory features (Table 2).

**Table 2.** Statistics of processed variants in *Cucumis sativus* var. hardwickii.

|  |  |
| --- | --- |
| Lines of input read | 6404727 |
| Variants processed | 2228224 |
| Variants filtered out | 0 |
| Novel / existing variants | - |
| Overlapped genes | 23777 |
| Overlapped transcripts | 23777 |
| Overlapped regulatory features | - |

#### 3.2.2 Variants classes

Single-nucleotide variants (SNV) are the most commonly present variants genome wide and detected with great value in present study. Total identified variants in present study with computational analysis are segregated mainly into four classes of genomic variants which includes 0.01% Sequence alteration (275), 5.94% Insertion (1,32,531), 6.37% Deletion (1,42,040) and 87.66% SNV (19,53,378) respectively (Table 3).

**Table 3.** Summary of predicted variant classes in *Cucumis sativus* var. hardwickii.

| Variant class | Count | % |
| --- | --- | --- |
| Sequence alteration | 275 | 0.01 |
| Insertion | 132,531 | 5.94 |
| Deletion | 142,040 | 6.37 |
| SNV | 1,953,378 | 87.66 |
| Total | 2228224 |  |

#### 3.2.3 All Consequences and most severe consequences of variants

Most severe consequences were detected in 21 classes with significant difference in number, highest for upstream gene variants i.e., 1,029,734 and lowest for Start retained variant i. e., 02 only. Some major classes are frame shift variants (1,114), inframe insertion (588), inframe deletion (712), missense variant (38,641), protein altering variant (29), synonymous variant (42,823), coding sequence variant (21), 5’ prime UTR variant (20,079), 3’ prime UTR variant (27,691), intron variant (3,04,952), downstream gene variant (3,66,513) and intergenic variant (3,87,161) were also detected for most severe consequences. Similarly, All consequences were detected in 21 classes with great value change, highest for upstream gene variants i.e., 1,547,272 and lowest for start retained variant (13) (Table 4).

**Table 4.**
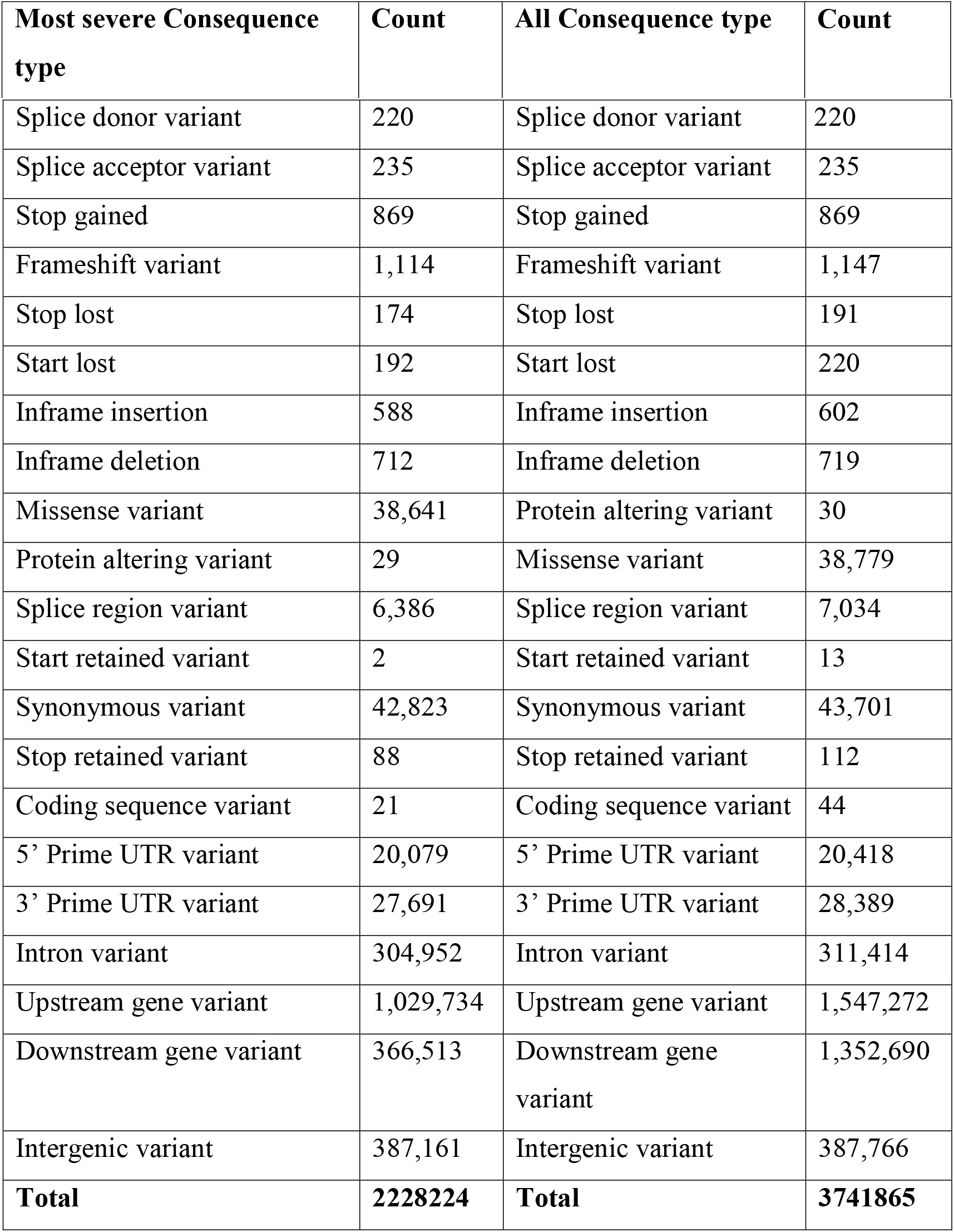
Most severe consequences and all consequences type data in *Cucumis sativus* var. hardwickii.

| Most severe Consequence type | Count | All Consequence type | Count |
| --- | --- | --- | --- |
| Splice donor variant | 220 | Splice donor variant | 220 |
| Splice acceptor variant | 235 | Splice acceptor variant | 235 |
| Stop gained | 869 | Stop gained | 869 |
| Frameshift variant | 1,114 | Frameshift variant | 1,147 |
| Stop lost | 174 | Stop lost | 191 |
| Start lost | 192 | Start lost | 220 |
| Inframe insertion | 588 | Inframe insertion | 602 |
| Inframe deletion | 712 | Inframe deletion | 719 |
| Missense variant | 38,641 | Protein altering variant | 30 |
| Protein altering variant | 29 | Missense variant | 38,779 |
| Splice region variant | 6,386 | Splice region variant | 7,034 |
| Start retained variant | 2 | Start retained variant | 13 |
| Synonymous variant | 42,823 | Synonymous variant | 43,701 |
| Stop retained variant | 88 | Stop retained variant | 112 |
| Coding sequence variant | 21 | Coding sequence variant | 44 |
| 5' Prime UTR variant | 20,079 | 5' Prime UTR variant | 20,418 |
| 3' Prime UTR variant | 27,691 | 3' Prime UTR variant | 28,389 |
| Intron variant | 304,952 | Intron variant | 311,414 |
| Upstream gene variant | 1,029,734 | Upstream gene variant | 1,547,272 |
| Downstream gene variant | 366,513 | Downstream gene variant | 1,352,690 |
| Intergenic variant | 387,161 | Intergenic variant | 387,766 |
| **Total** | **2228224** | **Total** | **3741865** |

#### 3.2.4 Coding consequences and variant location on chromosomes

A total of 86435 variants were recorded for Coding consequences (a variant that makes changes in coding sequence) and classified into 12 classes with variable count range and highly detected 50.55% (43,701) in Synonymous variant (A sequence variant changes one base of terminator codon but does not impact on encoded amino acids, the terminator remains), followed by 44.86% (38,779) Missense variant (A variant that make changes in one base of the transcript first codon) and least detected for 0.01% (13) Start retained variant class (Table 5). A total of 22,28,224 variants were detected by Ensembl VEP, although variants were identified in all chromosomes, the number of detected variants on chromosomes varied considerably. These variants distributed on all seven chromosomes of *Cucumis sativus* var hardwickii with high number 18.03% (401,920) of total detected variants on chromosome 3 followed by 15.84% (352,961) chromosome 5, 15.46% (344,544) chromosome 1, 14.72% (328,039) chromosome 6, 12.92% (287,924) chromosome 2, 12.13% (270,449) chromosome 4 while less 10.87% (242,387) on chromosome 7 (Table 6).

**Table 5.**
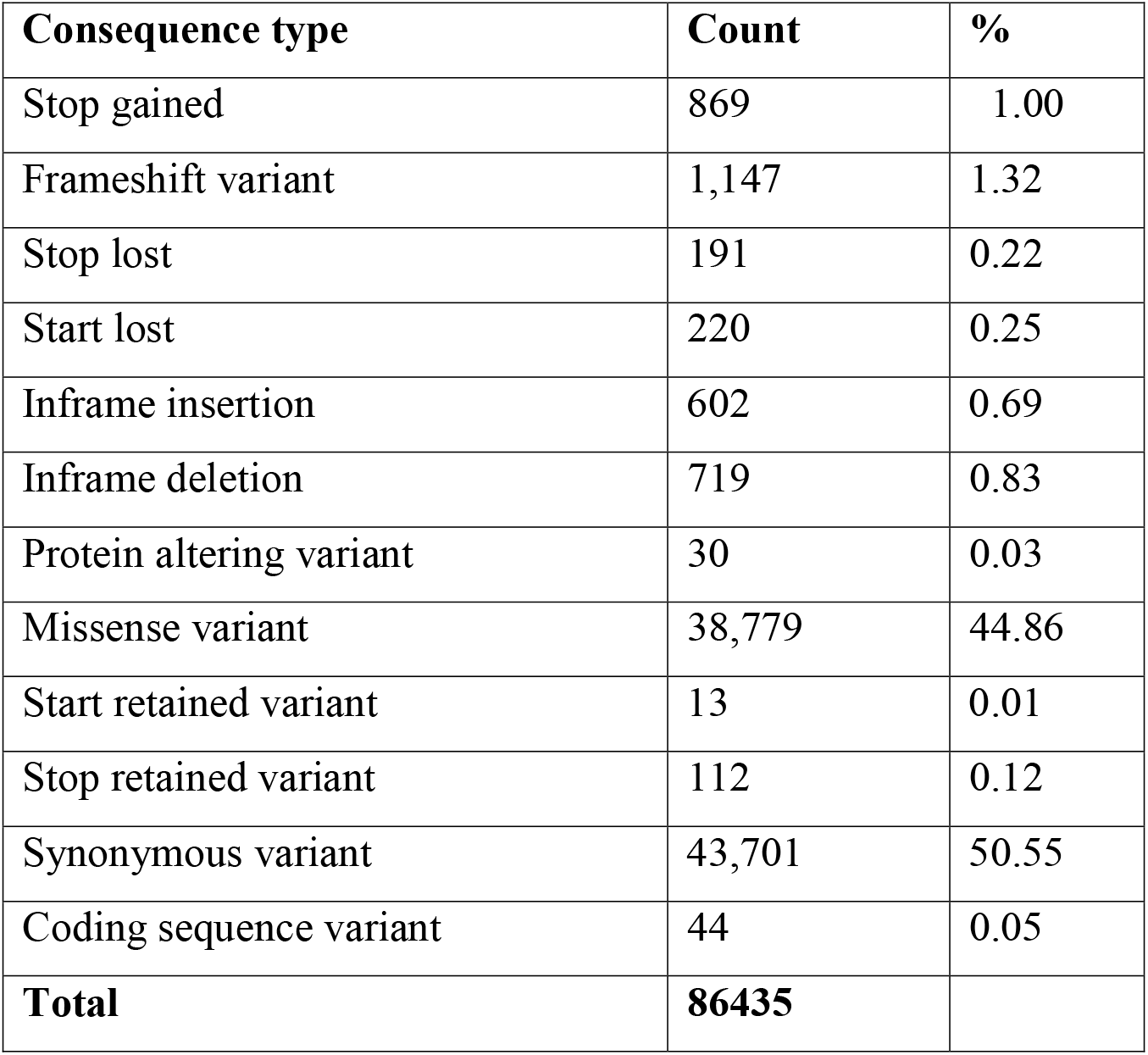
Coding consequences identified in *Cucumis sativus* var. hardwickii.

**Table 6.**
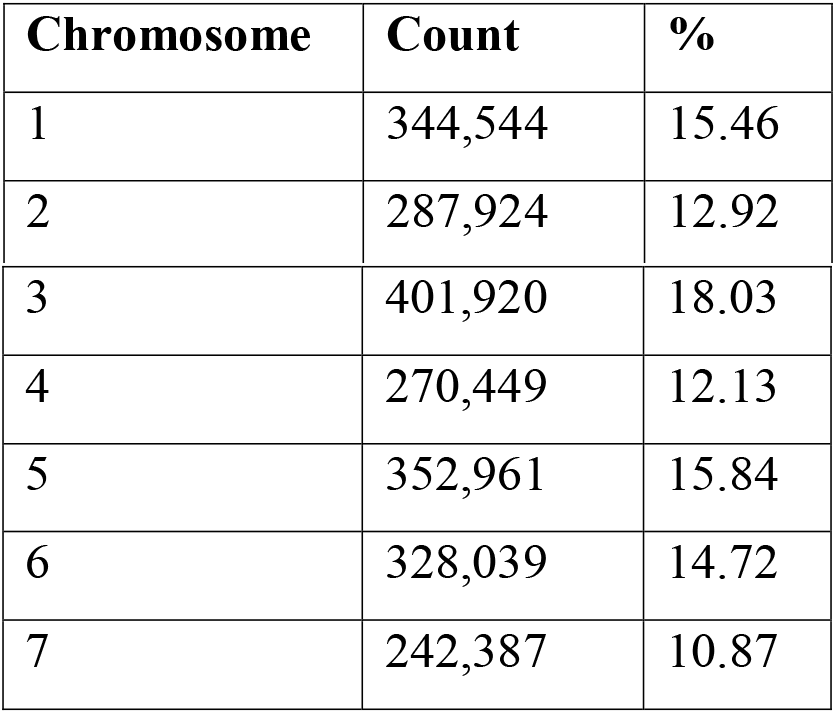
Chromosomal distribution of predicted variants in the *Cucumis sativus* var. hardwickii genome.

| Chromosome | Count | % |
| --- | --- | --- |
| 1 | 344,544 | 15.46 |
| 2 | 287,924 | 12.92 |
| 3 | 401,920 | 18.03 |
| 4 | 270,449 | 12.13 |
| 5 | 352,961 | 15.84 |
| 6 | 328,039 | 14.72 |
| 7 | 242,387 | 10.87 |

## 4. Discussion

Cucumber is a vegetable crop, the economic importance of which varies according to the region of the world. Cucumber is widely considered an excellent model for studying sex determination in monoecious plants (Pawełkowicz et al., 2019). Cucumber plant breeding can benefit from research into the phenotypic diversity found in cucumber genotypes. Superior genotypes for fruit yield and component production could be used to increase fruit yield, and they are also recommended for hybridization in cucumber cultivation (Golabadi et al., 2012). Next-generation sequencing (NGS) offers a new perspective on the genetic code, revealing new genes and regulatory sequences, as well as a wealth of molecular markers. When it comes to traits that are influenced by a large number of factors, expression studies provide breeders with a wealth of knowledge about the molecular basis of these traits (Pawekowicz et al., 2016). Advances in next-generation sequencing technology have facilitated the widespread use of SNPs in crops. Computational approaches dominate SNP discovery methods because increasing sequence awareness in public databases makes it easier to identify insightful SNPs computationally; however, complex genomes complicate SNP discovery significantly more, necessitating the use of alternative strategies for some crops. Plant genomes, in their entirety, have been shown to be repetitive (Mammadov et al., 2012).

Genome-wide sequencing analysis for Single Nucleotide Polymorphisms and Variants prediction, along with their genome-wide distribution, may help to improve our understanding of genetic diversity, evolutionary studies and superior economically important traits of crop plants (Cavagnaro et al., 2010).In this study, we conducted genome-wide SNPs identification and variants analysis from whole-genome sequencing data of *Cucumis sativus var* hardwickii. SNPs are very useful for studying genetic diversity because they can reveal the connections between different varieties. SNPs have been used for several years to measure diversity in specific genes or genomic regions, and the results are used to infer phylogenetic relationships between species (Morgil et al., 2020). The successful mapping of 92.9M raw reads with the reference genome revealed 19, 74,213 SNPs and 22, 28,224 variants genome-wide. SNPs allowed us to better understand genetic diversity and agricultural improvement, which will result in the enhancement of critical crop traits.

Variants calling in cucumber mapped sequencing data resulted in four variant classes with high number of SNV (87.66%) followed by deletion (6.37%), insertion (5.94%) and negligible for sequence alteration (0.01%). Here we recorded 21 variants classes for the most severe consequences with a total count of 2228224, highest for upstream gene variants (1,029,734), which located in 2 Kb 5’ of a gene followed by Intergenic variant found in intergenic regions (387161) and lowest for Start retained variant (02). Similarly, 21 variant classes for all consequences type with total number 3741865, highly detected for upstream gene variants, i.e., 1,547,27, followed by Downstream gene variant, i.e., 1352690 and lowest for Start retained variant, i.e., 13. Based on the variant location in the protein-coding and non-coding regions, including non-synonymous, missense, nonsense or Frameshift variants (Shameer et al., 2016).

Impact of variants for coding consequences category revealed high change is synonymous (50.55%) where one base of stop codon changed. Still, terminator remains and does not affect the encoding amino acid peptide, missense (44.86%), which change at least one base in start codon and affect the encoding protein, followed by frame shift (1.32%) which resulting in disruption of translating reading frame as insertion or deletion is not multiple of three and stop gained (1.00%) (Liu et al., 2019). The discovered Variants were distributed across all 7 chromosomes. Overall, 18.03% (401920) of the total discovered variants (2228224) were on chromosome 3, followed by 15.84% (352961) on chromosome 5 and at least 10.87% (242387) on chromosome 7. The above genome-wide analyzed SNPs and Variants may improve our understanding of genetic evolution and diversity in Cucumis sativus and significantly increase our knowledge about the variants’ association with genomic diversity and plants’ response to environmental factors.

## 5. Conclusion

Genome-wide sequencing data have become a powerful tool for exploring the genetic diversity and functional significance of SNP) and variants in various organisms. These molecular markers can reveal the evolutionary history, population structure, and adaptive responses of different species and cultivars. In this study, we performed a comprehensive analysis of SNPs and variants in the resequenced genome of Cucumis sativus var. hardwickii, a wild relative of cultivated cucumber. We detected a total of 1974213 SNPs and 2228224 variants across the seven chromosomes of C. sativus var. hardwickii, with chromosome 3 having the highest density and chromosome 7 having the lowest density. These genomic variations reflect the high level of genetic diversity and divergence between C. sativus var. hardwickii and other cucurbit species. Furthermore, we annotated the functional effects of these SNPs and variants on genes and regulatory elements involved in various biological processes related to plant growth, development, stress tolerance, and disease resistance. Our results provide valuable resources for future studies on the molecular basis and phenotypic consequences of genome-wide variation in C. sativus var. hardwickii and its potential applications for cucumber breeding.

